# CellStitch: 3D Cellular Anisotropic Image Segmentation via Optimal Transport

**DOI:** 10.1101/2023.06.19.545594

**Authors:** Yining Liu, Yinuo Jin, Elham Azizi, Andrew J. Blumberg

**Affiliations:** Department of Computer Science, Columbia University, New York, USA; Department of Biomedical Engineering, Columbia University, New York, USA; Department of Mathematics, Columbia University, New York, USA; Data Science Institute, Columbia University, New York, USA; Irving Institute for Cancer Dynamics, New York, USA

**Author notes:** These authors contributed equally to this work (Yining Liu, Yinuo Jin).

**Keywords:** Bioimaging, Optimal transport, 3D segmentation

## Abstract

**Background:** Spatial mapping of transcriptional states provides valuable biological insights into cellular functions and interactions in the context of the tissue. Accurate 3D cell segmentation is a critical step in the analysis of this data towards understanding diseases and normal development *in situ*. Current approaches designed to automate 3D segmentation include stitching masks along one dimension, training a 3D neural network architecture from scratch, and reconstructing a 3D volume from 2D segmentations on all dimensions. However, the applicability of existing methods is hampered by inaccurate segmentations along the non-stitching dimensions, the lack of high-quality diverse 3D training data, and inhomogeneity among different dimensions; as a result, they have not been widely used in practice.

**Methods:** To address these challenges, we formulate the problem of finding cell correspondence across layers with a novel optimal transport (OT) approach. We propose CellStitch, a flexible pipeline that segments cells from 3D images without requiring large amounts of 3D training data. We further extend our method to interpolate internal slices from highly anisotropic cell images to recover isotropic cell morphology.

**Results:** We evaluated the performance of CellStitch through eight 3D plant microscopic datasets with diverse anisotropic levels and cell shapes. CellStitch substantially outperforms the state-of-the art methods on anisotropic images, and achieves comparable segmentation quality against competing methods in isotropic setting. We benchmarked and reported 3D segmentation results of all the methods with instance-level precision, recall and average precision (AP) metrics.

**Conclusion:** The proposed OT-based 3D segmentation pipeline outperformed the existing state-of-the-art methods on different datasets with nonzero anisotropy, providing high fidelity recovery of 3D cell morphology from microscopic images.

## 1 Introduction

Spatial profiling of transcriptional states enables researchers to understand how the organization of cells influences function by providing spatial context to single cells. Indeed, spatially resolved transcriptomics was crowned as the Method of the Year in 2020 due to the valuable biological insights it provides [1]. As a first step in the pipeline, segmentation defines cell boundaries from fluorescent staining signals, assisting the assignment of RNA amplicons to reconstruct a gene-cell matrix; hence it plays a crucial role in characterizing cell types, their morphology, and location in the context of their microenvironment [2]. Segmentation is also a critical step in the analysis of multiplexed imaging of proteins [3, 4].

Deep learning has been successful at 2D cell segmentation, leveraging the availability of diverse labeled 2D training data and specialized deep learning architectures such as U-Net [5]. Trained on labeled 2D images, 2D cell segmentation pipelines such as Mesmer [6], StarDist [7], and Cellpose [8] can segment cells from 2D images with minimal human supervision and achieve expert-level accuracy. On the other hand, due to the lack of large and diverse 3D training datasets, approaches that utilize 3D neural networks [9–12] do not have comparable accuracy and generalizability, particularly in complex or dense tissues comprised of heterogeneous and abnormally shaped cells such as cancer cells. Additionally, these methods often lead to the over-segmentation of cells or noisy masks, thus impacting downstream analyses. As 3D transcriptomics studies increase in scale and computational cost [13], the need for a robust, generalizable, and user-friendly 3D cellular instance segmentation pipeline has become increasingly urgent [10]. To leverage the accuracy of 2D segmentation, [8] performs 2D segmentation layer by layer along a stitching direction and declares two cell slices as coming from the same cell if their overlap exceeds a predefined threshold. However, when cells are stacked roughly on top of each other along the stitching direction, the resulting masks are inaccurate along the non-stitching directions. Alternative approaches have been proposed to segment cell slices on different projections of the images with subsequent 3D reconstruction [8, 14], but the performance of such methods often suffers because of *anisotropy* in the microscopic images, i.e., inhomogeneity of image resolution among different dimensions due to experimental constraints. For instance, the thickness of tissue resections can be determined by imaging technology, tissue availability, and ensuring preserved tissue architecture, hence leading to higher resolution in the imaging (*X* and *Y*) dimensions than along the slicing (*Z*) dimension for softer tissues.

Here we present CellStitch, a pipeline that applies optimal transport to segment and reconstruct cells from 3D images without requiring large 3D training datasets. Optimal transport studies the best way of transforming a source distribution into a target distribution [15]; it is a natural way to pose matching problems and hence has been widely studied in mathematics, economics, and statistics. Recently, optimal transport has also been applied in the computer vision and machine learning communities because of the computational efficiency of its relaxation [16].

CellStitch focuses on stitching 2D masks along one dimension. Hence, CellStitch does not require end-to-end training of any 3D network. Additionally, in contrast to a commonly-used stitching method [8] that fails to incorporate crucial information from the other directions, CellStitch uses optimal transport to trace cells across image layers and then uses segmentations from the other two directions to guide the declaration of new cells. In particular, to find correspondence between cells in adjacent layers, we model the layers as certain associated discrete distributions and obtain the optimal correspondence of cells based on the optimal transport plan. The CellStitch framework is illustrated in Fig. [1].

To summarize, the main contributions of our work are as follows:

- We formulate the problem of finding correspondence between cells from adjacent layers in terms of optimal transport using a novel cost matrix. In particular, we highlight the importance of using pairwise cell overlaps to compute the cost matrix for the given task instead of the widely-used pairwise distances.
- We design a framework to segment cells from 3D images that does not require end-to-end training of 3D networks. This is achieved by using the optimal transport plan to infer cell correspondence across different layers, and leveraging 2D segmentations from the other projections to resolve the stitching ambiguity of the matched cell slices.
- We further provide a novel interpolation framework based on optimal transport to interpolate adjacent layers; the interpolation extension can be used to reduce the anisotropy of images.
- We compare our 2D-based framework against the state-of-the-art 2.5D-based and 3D-based frameworks, and observe that CellStitch consistently performs better across datasets with anisotropy and is comparable to the best on isotropic datasets with spherical cells.

## 2 Related Work

With the recent emergence of imaging-based spatial transcriptomics and proteomics platforms [17, 18], accurate quantification of cell boundaries becomes increasingly essential. In recent years, the state-of-the-art deep convolutional neural networks (CNNs) based on U-Net [5] or ResNet backbones [19] have enabled successful 2D and 3D segmentation across various medical imaging domains [20, 21]. The increasing development of multiplexed 3D imaging technologies has driven higher demand for volumetric 3D segmentation.

To our knowledge, there are three major approaches for 3D instance segmentation: 3D end-to-end training, 2D segmentation layer by layer followed by stitching, and 2.5D projections followed by 3D reconstruction. The direct 3D approaches, such as PlantSeg [12] and 3DCellSeg [10], rely on 3D U-Net variants trained upon 3D annotated datasets. The 3D models enable smooth boundary predictions by incorporating contextual information from nearby layers in all directions, but suffer from high computational cost, lack of generalizability, and in some cases specific convex morphology assumptions [11]. For example, [10] reports over 10% performance drop in Jaccard Index (JI) and Dice Similarity Coefficient (DSC) when using test data collected using a different type of confocal laser scanning microscopy from the one used in their training dataset.

Meanwhile, the latest 2D segmentation pipelines [6, 8] have shown robust performance aided by diverse annotated data modalities and 2D models along *xy* planes. The final 3D reconstruction is then performed by stitching the output masks along the *Z*-axis with heuristic instance mapping methods such as Intersection-over-Union (IoU). Their performance is sensitive to the empirical stitching threshold; optimal threshold values are data-dependent and hard to determine *a priori*. Moreover, the existing 2D-based approaches fail to resolve the stitching ambiguity imposed by the arrangement of cells. Empirically, we observe a nontrivial number of pairs of cell slices that have significant overlap but are from distinct cells, because distinct cells are stacked on top of each other; as a result, existing 2D-based approaches tend to produce under-segmented masks on the projections along the *X*- and *Y* -axes.

The 2.5D segmentation approaches [8, 14] attempt to combine the advantages of 2D and 3D, utilizing contextual layer awareness with efficient, transferable models. For instance, Cellpose3D [8] trains models to predict the flow vectors for each pixel; in order to obtain the 3D flow vectors, Cellpose3D averages the 2D flow vectors along the *XY, XZ* and *Y Z* directions. Nevertheless, the substantial inhomogeneous sampling ratios between *XY* plane and *Z*-axis introduce noise to the segmentation pipeline, leading to over-segmentation on highly anisotropic images.

Recently, algorithms based on ideas from optimal transport theory have begun to find applications in biology [22, 23]. In particular, PASTE [24] performs pairwise spot alignment across adjacent Visium layers [25], and SCOTT [26] designs a shape-location combined system for cell tracking in 2D microscopy videos. Other applications of optimal transport have addressed 2D and 3D image registrations such as retinal fundus alignment [27, 28]. However, those methods cannot be easily applied to fluorescent images with dense, cluttered cells, because they mainly focus on spot or pixel-level alignment with single or at most a few objects of interest. Therefore, to overcome the limitations of direct 3D segmentation, CellStitch applies optimal transport to align cell objects generated from any given well-trained 2D segmentation model along the *Z* direction in order to reconstruct final 3D segmentations.

## 3 Methods

### 3.1 Background

For discrete measures, the Kantorovich formulation of the optimal transport problem is as follows: given two discrete measures *P* : [*m*] → ℝ and *P*^′^ : [*n*] → ℝ, a transport plan 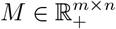 is a product distribution on [*m*] ×[*n*] that has *P* and *P* ^′^ as marginals; in other words, *M*_*x,y*_ describes the amount of mass in bin *x* that flows to bin *y*. Additionally, given a cost matrix *C* ∈ ℝ ^m×n^ such that *C*_*x,y*_ describes the price associated with moving a unit of mass from bin *x* to bin *y*, the Kantorovich optimal transport solves for

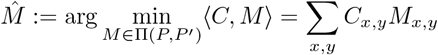

where

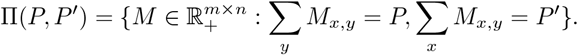

The choice of the cost matrix *C* determines the properties of the optimal transport problem such as the uniqueness of solutions and the computational cost of solving the optimization problem. A popular choice to compute the cost matrix is the *l*_*p*_ distance, where *l*_*p*_(*x, y*) := ‖*x* – *y*‖^*p*^. In this case, optimal transport induces a distance on the space of distributions, referred to as the *p*-Wasserstein distance. In particular, the 2-Wasserstein distance is obtained by using the Euclidean distance as the cost matrix; it has gained popularity in the computer vision community for performing interpolation, color transfer, and geometry processing [16, 29].

### 3.2 Optimal Transport for Stitching Adjacent Layers

Suppose there are *m* + 1 labels in layer *z* and *n* + 1 labels in layer *z* + 1 along *Z*-axis, where the label 0 is reserved for background pixels and any label *l >* 0 represents a cell label; the goal is to relabel pixels in layer *z* + 1 so that the pixels corresponding to the same cell have the same label in the two layers.

In terms of optimal transport, we want to find the optimal transport plan between the two discrete distributions *P* : [*m* + 1] → ℝ and *P* ^′^ : [*n* + 1] → ℝ where

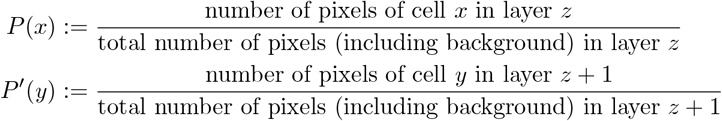

i.e. the distributions are proportions of label masses.

The cost matrix *C* ∈ ℝ ^(*m*+1)×(*n*+1)^ is defined as

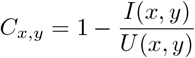

where

- *I*(*x, y*) is the number of pixels in the intersection of cell *x* in layer *z* and cell *y* in layer *z* + 1;
- *U* (*x, y*) is the number of pixels in the union of cell *x* in layer *z* and cell *y* in layer *z* + 1.

The solved transport plan 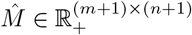 tells us the optimal way of moving labels in layer *z* to labels in layer *z* + 1.

Note that, unlike the natural approach of defining the cost matrix in terms of Euclidean distances, using pairwise distances between cell centroids would not give us an optimal assignment in this case. Consider the example shown in Fig. 2: the green and red cells would be incorrectly matched if we were to use the pairwise distance between the centroids as the cost matrix.

**Figure 1:**
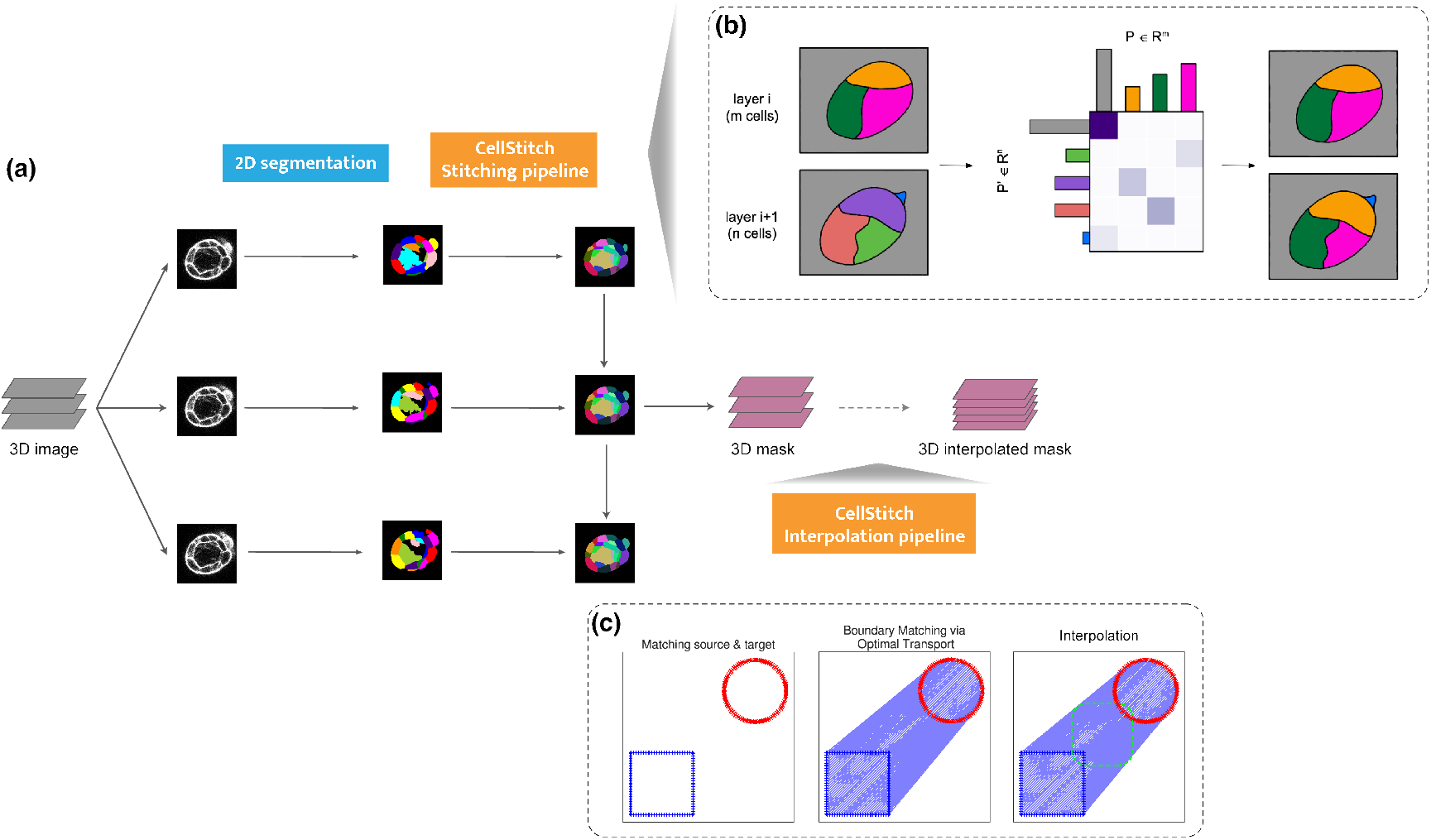
Overview of CellStitch framework. (a) CellStitch consists of a 3-step pipeline to reconstruct 3D cells: backbone 2D segmentation along *Z*-axis direction, stitching module, and an optional interpolation module. (b) Stitching pipeline: CellStitch first computes the source and target distributions based on the cell masses on given adjacent layers, and a cost matrix based on the pairwise overlap. The solved optimal transport plan is then used to deduce the optimal correspondence of cells across the adjacent layers. Finally, it reassigns instance labels of cell slices to enforce labeling consistency across all layers and creates new labels when new cells emerge based on the segmentations masks on the other two projections. (c) Interpolation pipeline: After finding optimal instance stitching, CellStitch leverages pixel-wise boundary matching followed by morphology interpolation to predict the internal layers between adjacent slices.

**Figure 2:**
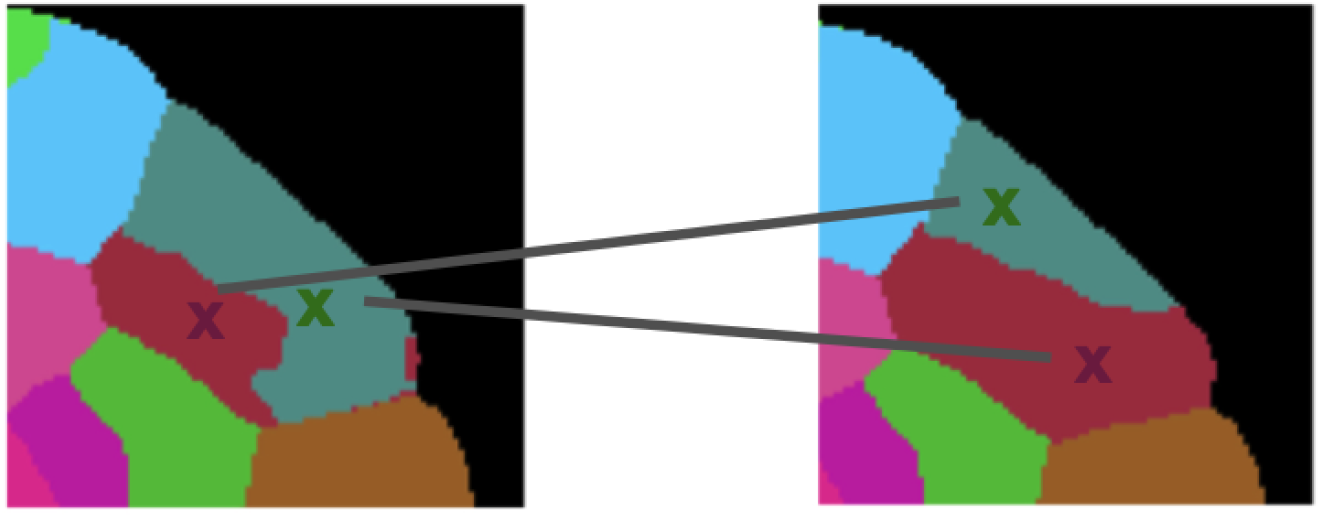
Poor correspondence of cell centroids. Using the pairwise distances between cell centroids would result in mismatched cells; as a result, this motivates the use of the union over intersection to compute the cost matrix. Here, the color code is based on the ground truth correspondence while the edges highlight the misassignments based on cell centroids. The image is taken from cell labels from the ovules dataset [30].

### 3.3 CellStitch Stitching Algorithm

We now describe how we use the optimal transport plan 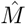 computed in Sect. 3.2 to relabel cells in layer *z* + 1 consistently with the labels in layer *z*.

For each cell, *x* in layer *z*, notice that *ŷ* := arg max_*y*_ *M*_*x,y*_ is the cell in layer *z* + 1 such that cell *x* most likely moves to. Define the cell tracing function *T*_*M*_ : [*m* + 1] → [*n* + 1] induced by the transport plan *M* as

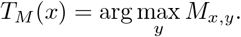

If 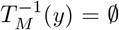 (i.e. no cells in layer *z* to trace the cell *y*), we declare *y* as a new cell and assign it a new label *l* ∉ [*n* + 1]. Note that *T*_*M*_ might not be an injection; for a given cell in layer *z* + 1, there can be more than one cell in layer *z* that traces to *y*. As a result, for each cell *y* ∈ [*n* + 1], we match it to the cell tracing to it in layer *z* with the least cost:

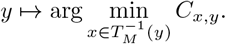

Notice that the matched cell slices should not always be assigned to the same labels, because they might come from distinct cells if the two cells are stacked on top of each other in the *Z*-axis direction. In order to resolve the ambiguity imposed by the placement of cells, we use a voting scheme to decide whether two OT-matched cell slices will be stitched or not. Each cell pixel in the matched cell in layer *z* + 1 gets two votes to reject stitching. For a given pixel, we use the 2D segmentation masks (*Y Z*-masks and *XZ*-masks) in the projections to *X*- and *Y* -axis to determine whether they vote for rejecting stitching or not. In particular, a cell pixel will use one rejection vote if it gets assigned different cell labels in the *Y Z*-masks (similarly for the *XZ*-masks). At the end, the two cell slices will only be stitched (i.e. assigned as the same cell labels) if the proportion of rejection votes is smaller than a user-defined threshold. Algorithm 1 summarizes the stitching algorithm. In practice, we find that the choice of the stitching direction has little effect on the final segmentation results; open source code implementing the stitching algorithm from the top to the bottom layer is available at https://github.com/imyiningliu/cellstitch.

### 3.4 CellStitch Interpolation Algorithm

Due to cost and technology constraints of 3D imaging as well as physical characteristics of tissues, the *Z*-direction resolution is oftentimes lower than that of the *XY* -plane, introducing anisotropy to the 3D cell images. However, since cells are locally cylindrical, one could hope to decrease the anisotropy of the dataset by predicting the missing layers between adjacent slices.

We now present a pixel-level OT interpolation application of our pipeline after instance-level stitching. Given matched cell *c*_*s*_ in layer *z* and cell *c*_*t*_ in layer *z* + 1, we aim to reconstruct the shape of cells {*c*_1_, …, *c*_*N*−1_} located in equally distributed internal layers 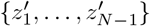 between *z* and *z* + 1, assuming the anisotropy of the original images is *N*.

Here we reformulate the problem as a boundary-matching task between the contour pixels across the source and target cells, and use Wasserstein interpolation [29] to infer the internal layers (Algorithm 2). In particular, in order to compute a geometry-aware average between two cell slices from adjacent layers, we first compute the optimal transport plan that achieves the 2-Wasserstein distance between the two uniform distributions on the cell boundaries. The optimal transport plan gives a partial matching between the pixels in the source cell boundary and the pixels in the target cell boundary. In order to interpolate the two slices, we compute the weighted average of the coordinates between each partially matched source and target pixel pairs matched target pixels locations, where the weights are determined by the transport plan, and further predicted the corresponding cell instances by filling the interpolated boundaries (Fig. 3). In practice, the interpolation is implemented via vectorization to avoid redundant inner-layer for loops (Algorithm 2). Open source code implementing the interpolation algorithm is also available at https://github.com/imyiningliu/cellstitch.

**Figure 3:**
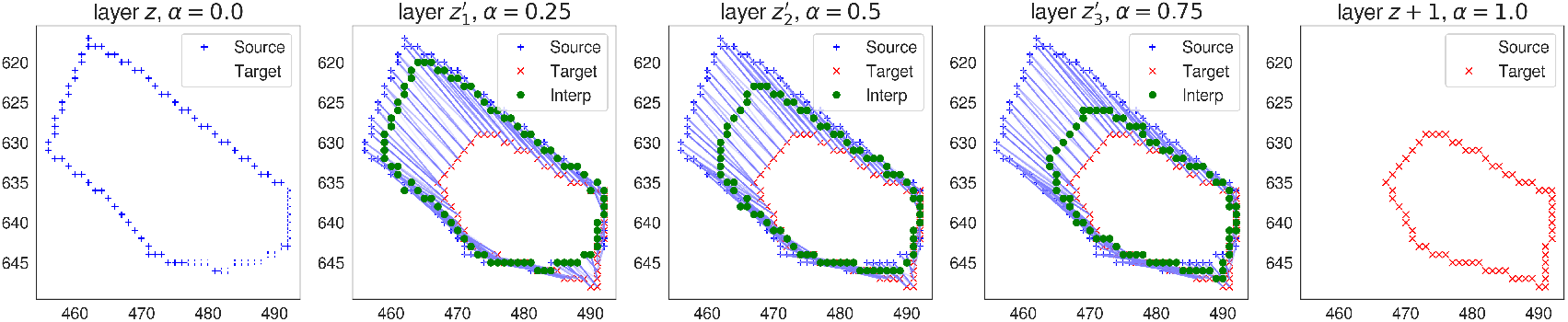
CellStitch interpolation diagram. Morphology interpolation of internal layers from an example pair of matched ovules cells in adjacent layers (anisotropy=4).

#### Algorithm 1 CellStitch: Stitching Algorithm

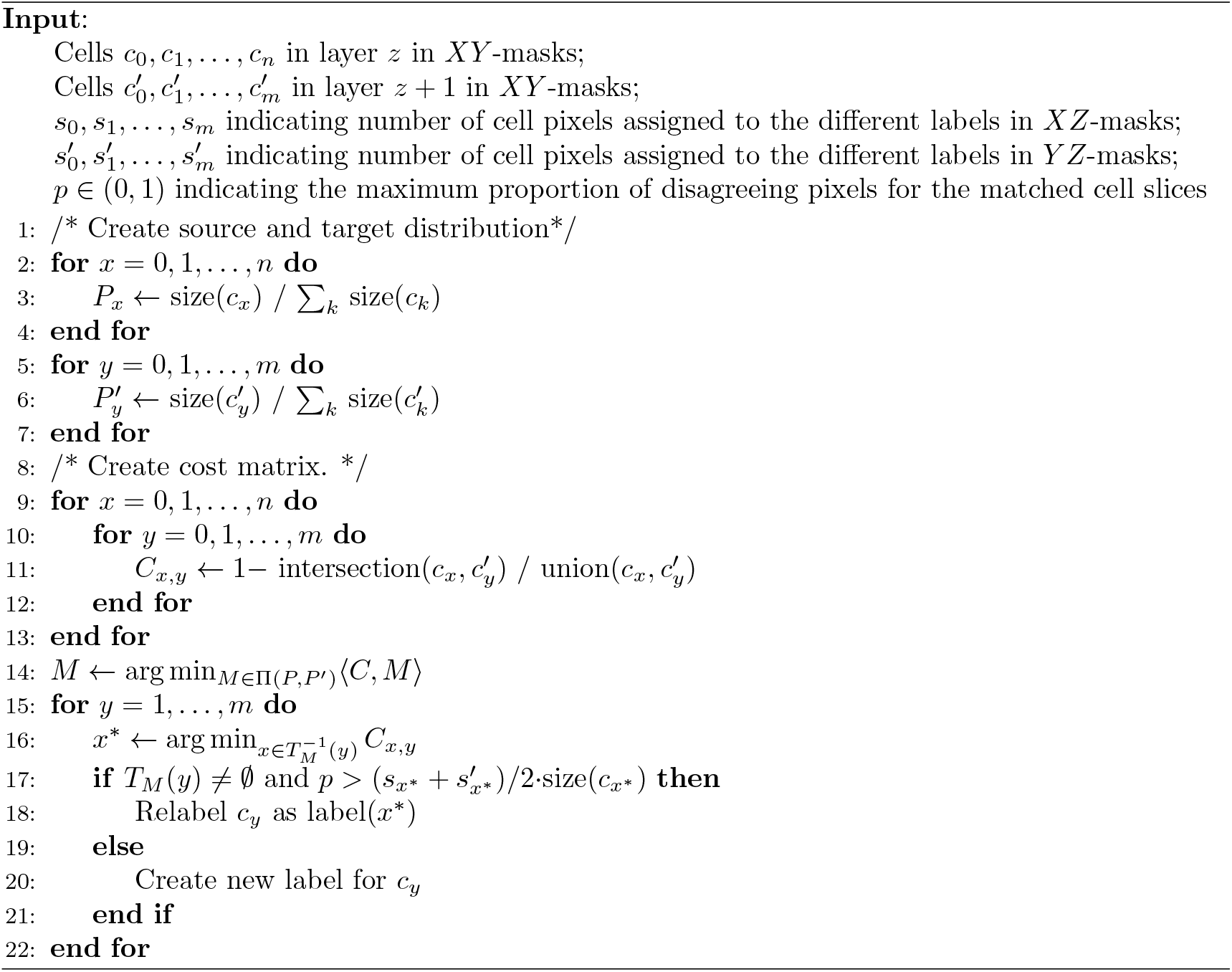

#### Algorithm 2 CellStitch: Interpolation Algorithm

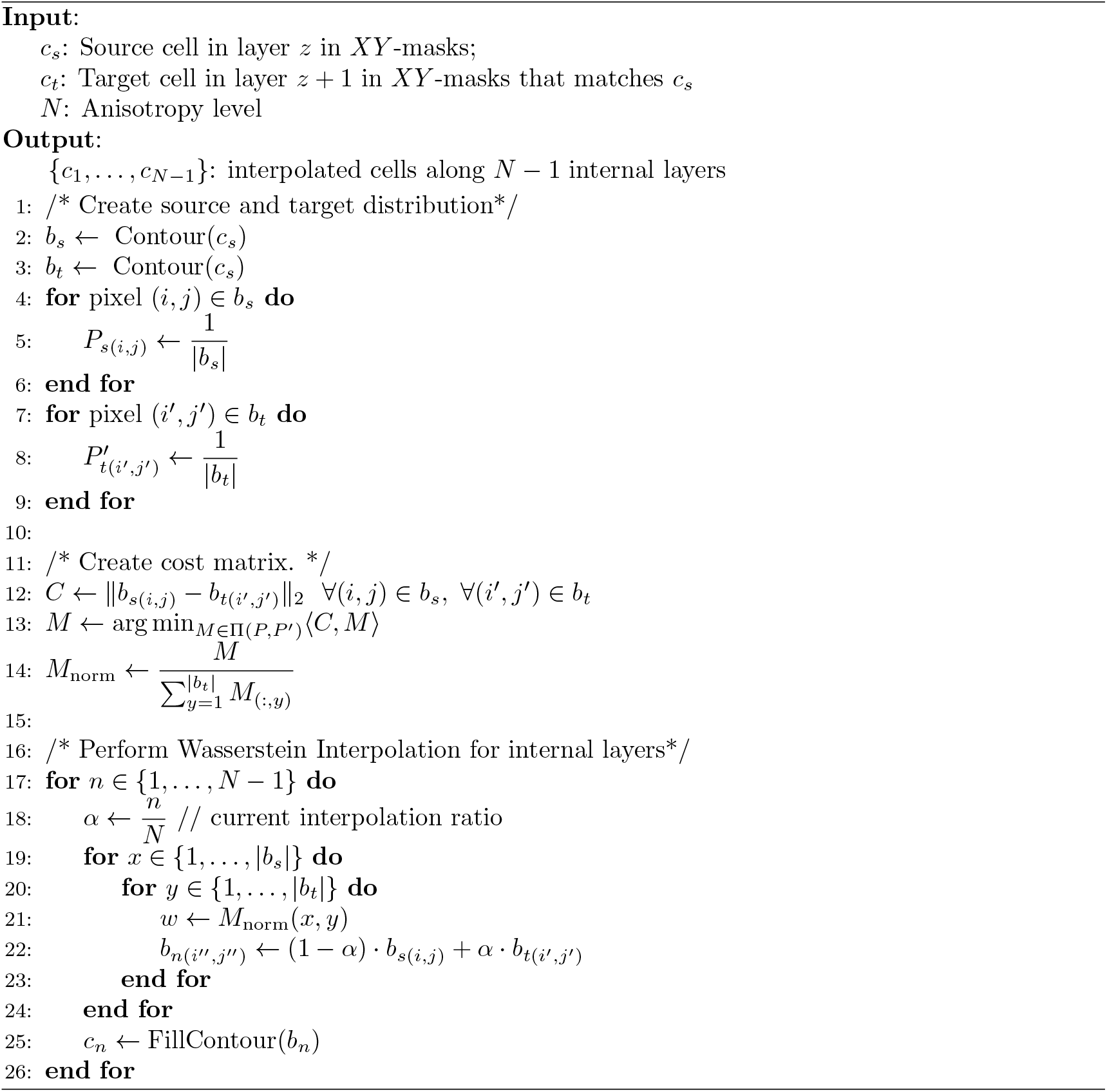

## 4 Results

### 4.1 Datasets

We evaluated CellStitch on eight publicly available *Arabidopsis thaliana* datasets with ground-truth segmentation labels (Table 1). The first dataset (ovules) [30] contains 31 anisotropic images of ovules cells at all developmental stages using confocal laser scanning microscopy (anisotropy = 4); the ovules dataset was used to benchmark CellStitch’s stitching and pipeline performance. The second dataset (ATAS) [31] contains 125 isotropic images of apical stem cells; the ATAS dataset was used to evaluate CellStitch’s performance at increasing anisotropy levels. Finally, in order to evaluate CellStitch’s performance under a realistic setting where the anisotropy of the dataset is unknown, we further generated six additional datasets by subsampling 600 images from six different plant organs from the Arabidopsis 3D Digital Tissue Atlas (https://osf.io/fzr56).

**Table 1:** Summary of benchmarking datasets

| Dataset | Number of images | Average image size ($Z \times Y \times X$ pixels) | Resolution ($Z \times Y \times X \mu m$) |
| --- | --- | --- | --- |
| Ovules | 31 | $317 \times 910 \times 949$ | $0.240 \times 0.063 \times 0.063$ |
| Apical Stem | 125 | $197 \times 509 \times 509$ | $0.26 \times 0.22 \times 0.22$ |
| Anther | 100 | $20 \times 224 \times 224$ | Unknown |
| Filament | 100 | $20 \times 224 \times 224$ | Unknown |
| Leaf | 100 | $20 \times 224 \times 224$ | Unknown |
| Pedicle | 100 | $20 \times 224 \times 224$ | Unknown |
| Sepal | 100 | $20 \times 224 \times 224$ | Unknown |
| Valve | 100 | $20 \times 224 \times 224$ | Unknown |

### 4.2 Evaluation Metrics and Benchmark Methods

We benchmarked our results with the state-of-the-art pipeline from each of the three classes of current deep learning-based 3D segmentation pipelines:

- 2D Cellpose with stitching via heuristic threshold (2D-based) [8],
- Cellpose3D (2.5D-based) [8],
- PlantSeg’s pretrained `confocal_unet_bce_dice ds1x` model (3D-based) [12].

In order to evaluate the segmentation accuracy, we first matched the cells in the segmentation mask and the ground truth label if the two cells have an intersection over union greater than 0.5. Then, we computed the precision, recall, and average precision as

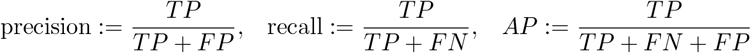

where *TP* is the number of matched cells, *FP* is the number of unmatched cells in the segmentation mask, and *FN* is the number of unmatched cells in the ground truth labels.

### 4.3 Experimental Results

We first evaluate CellStitch’s segmentation performance on the ovules dataset [30] under two settings:

- low anisotropy (anisotropy = 4): original data;
- high anisotropy (anisotropy = 8): sparsifying the *Z*-dimension by removing every other layer.

To compare the performance between the 2D-based, 2.5D-based, and 3D-based methods, we used the same training data (22 training images, 2 validation images, 7 test images) that was used to train PlantSeg’s `‘confocal_unet_bce_dice_ds1x’` model to train a Cellpose 2D segmentation model for 100 epochs with learning rate 0.2 and batch size 8; the trained model was then used as a backbone to generate 2D masks later used for CellStitch’s stitching and Cellpose3D’s reconstruction. We see that Cellpose3D suffers the most from increased anisotropy in the raw data, and that CellStitch consistently produces the best segmentation masks for both low and high anisotropy settings (Table 2, Fig. 4, 5). Due to the observation that PlantSeg’s performance is not on par with the rest of the methods, as well as the lack of instructions on how to train PlantSeg models on additional datasets, we decided to focus on comparing the performance of CellStitch against Cellpose2D and Cellpose3D for the rest of the evaluation.

**Table 2:** Performance benchmarks on the ovules dataset; the **best** performance (within 0.03) is in bold. The method that achieves the best performance under the majority of the metric is marked with ^*^.

| Setting | Method | Precision | Recall | AP |
| --- | --- | --- | --- | --- |
| Low Anisotropy | CellStitch* | **$0.64 \pm 0.08$** | $0.64 \pm 0.14$ | **$0.48 \pm 0.11$** |
| | Cellpose2D | $0.42 \pm 0.07$ | $0.57 \pm 0.09$ | $0.31 \pm 0.05$ |
| | Cellpose3D | $0.45 \pm 0.20$ | **$0.84 \pm 0.11$** | $0.42 \pm 0.20$ |
| | PlantSeg | $0.09 \pm 0.04$ | $0.17 \pm 0.05$ | $0.06 \pm 0.02$ |
| High Anisotropy | CellStitch* | **$0.66 \pm 0.07$** | **$0.52 \pm 0.10$** | **$0.41 \pm 0.08$** |
| | Cellpose2D | $0.48 \pm 0.05$ | **$0.54 \pm 0.09$** | $0.34 \pm 0.05$ |
| | Cellpose3D | $0.16 \pm 0.16$ | **$0.52 \pm 0.28$** | $0.15 \pm 0.14$ |
| | PlantSeg | $0.17 \pm 0.06$ | $0.26 \pm 0.04$ | $0.12 \pm 0.04$ |

**Figure 4:**
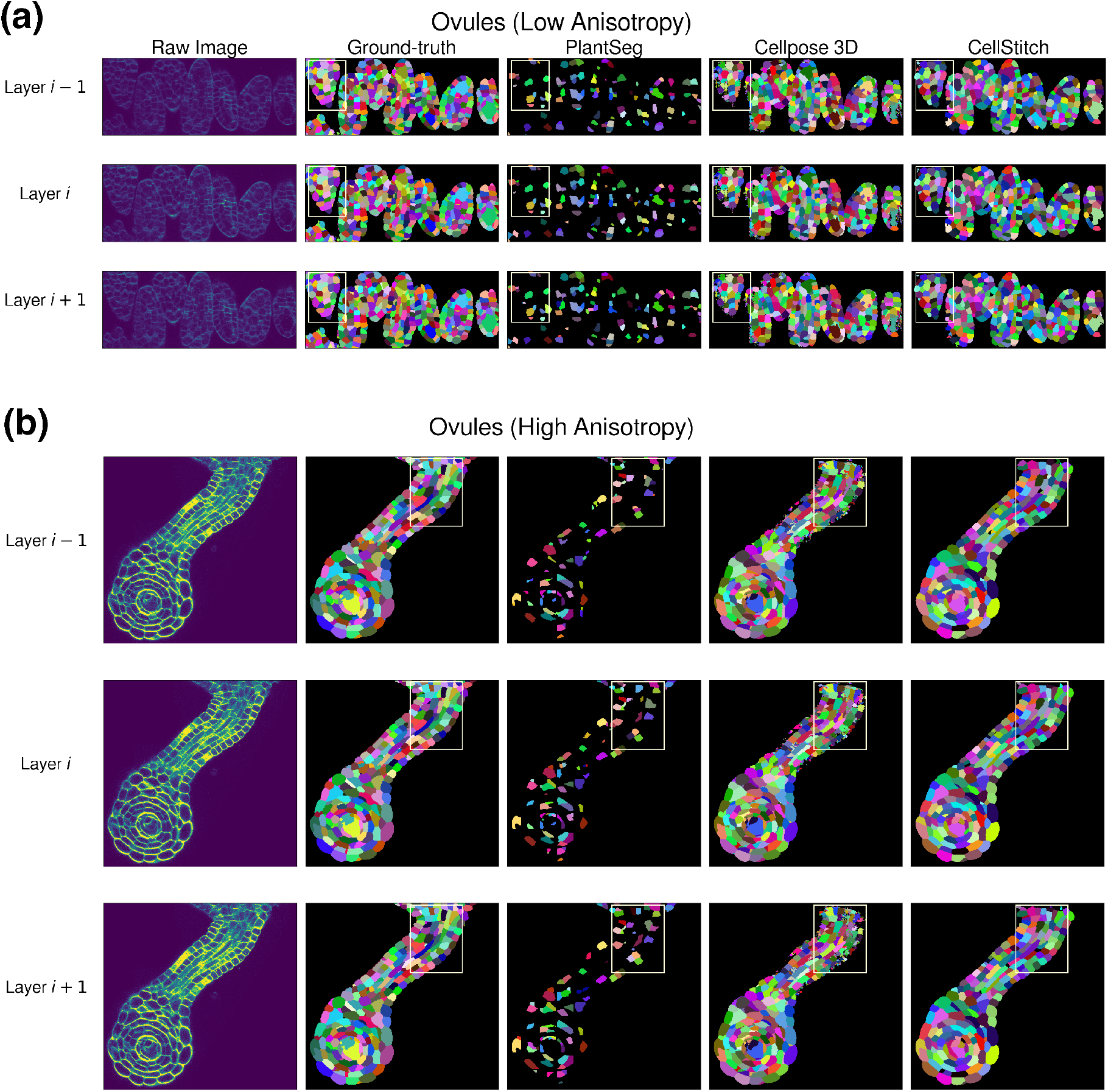
Qualitative pipeline benchmark results on consecutive Ovules slices. **(a) - (b)** Example low / high anisotropy confocal laser scanning microscopic images and ground-truth labels, along with PlantSeg, Cellpose3D, and CellStitch predictions are shown.

**Figure 5:**
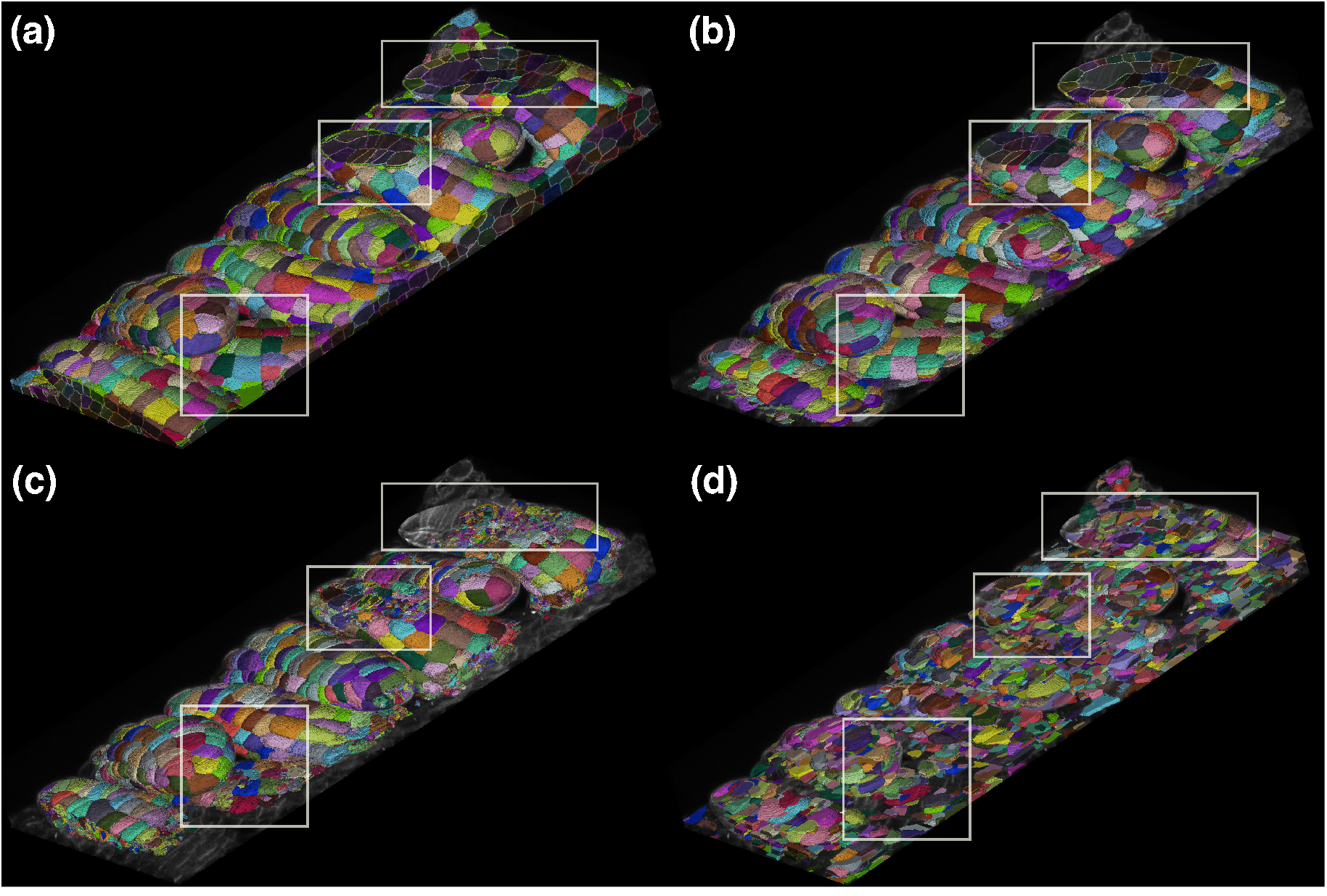
3D Visualization of segmentations. Example results of **(a)** Ground-truth, **(b)** CellStitch, **(c)** Cellpose3D, and **(d)** PlantSeg, overlaid with the raw Ovule images. Both Cellpose3D and PlantSeg tend to oversegment the cells (highlighted bounding boxes).

Then, we quantified the interpolation quality of CellStitch by reconstructing low anisotropy instances from high anisotropy images. We predicted the high anisotropy masks and further upsampled to the original *Z*-direction resolution. The interpolated masks achieved a mean average precision of 0.41; the results demonstrate that high anisotropy images achieved segmentations close to the original low anisotropy predictions of CellStitch, and both outperformed the baseline comparisons (Table 2).

Next, we used the *Arabidopsis thaliana apical* stem cells (ATAS) dataset [31] containing 125 isotropic images under 0.22*μm* × 0.22*μm* × 0.26*μm* resolution to further compare the performance of CellStitch and Cellpose; since the images in the ATAS dataset are isotropic, we are able to explore how the performance of CellStitch changes with increasing anisotropy. We introduced anisotropy in the ATAS dataset by subsampling across the *Z*-layers in order to further test the robustness of CellStitch and Cellpose3D against anisotropy in the dataset. We found that Cellpose3D achieves higher average precision when there is no anisotropy in the dataset, whereas Cellpose2D does not achieve comparable performance on isotropic dataset due to its ignorance of the other two directions. Additionally, similar to the results on the ovules dataset, the average precision of Cellpose3D significantly dropped under the high anisotropy setting. The quantitative results are presented in Table 3.

**Table 3:** Performance on the ATAS dataset; the **best** performance (within 0.03 is in bold. The method that achieves the best performance under the majority of the metric is marked with ^*^.

| Setting | Method | Precision | Recall | AP |
| --- | --- | --- | --- | --- |
| Anisotropy = 0 | CellStitch | $0.79 \pm 0.08$ | $0.77 \pm 0.04$ | $0.64 \pm 0.07$ |
| | Cellpose2D | $0.61 \pm 0.10$ | $0.69 \pm 0.04$ | $0.48 \pm 0.07$ |
|  | Cellpose3D* | **$0.87 \pm 0.17$** | **$0.98 \pm 0.01$** | **$0.85 \pm 0.17$** |
| Anisotropy = 5 | CellStitch* | **$0.81 \pm 0.04$** | **$0.58 \pm 0.05$** | **$0.51 \pm 0.05$** |
| | Cellpose2D | $0.67 \pm 0.04$ | **$0.61 \pm 0.05$** | $0.47 \pm 0.04$ |
| | Cellpose3D | $0.40 \pm 0.09$ | **$0.59 \pm 0.11$** | $0.32 \pm 0.08$ |
| Anisotropy = 10 | CellStitch* | **$0.75 \pm 0.04$** | $0.46 \pm 0.06$ | **$0.40 \pm 0.05$** |
| | Cellpose2D* | $0.64 \pm 0.04$ | **$0.55 \pm 0.05$** | **$0.42 \pm 0.05$** |
| | Cellpose3D | $0.30 \pm 0.09$ | $0.38 \pm 0.11$ | $0.20 \pm 0.08$ |

Finally, to test CellStitch’s performance under the practical setting, where the anisotropy of the datasets might be unknown, we generated six datasets by sampling manually annotated images from the Arabidopsis 3D Digital Tissue Atlas (https://osf.io/fzr56). For each of the six datasets, we trained Cellpose’s 2D segmentation model for 100 epoches with learning rate 0.2 and batch size 8 under a 7-3 train-test split as the backbone to further generate 3D segmentation results for CellStitch and Cellpose3D. We see in Table 4 that CellStitch is consistently the best performing method(s) on all datasets. Furthermore, CellStitch is also able to discover more morphology diversity, whereas Cellpose3D tends to over-segment non-spherical cells (see Fig. 6).

**Table 4:** Performance on Arabidopsis 3D Digital Tissue Atlas; the **best** performance (within 0.03) is in bold. The method that achieves the best performance under the majority of the metric is marked with ^*^.

| Dataset | Method | Precision | Recall | AP |
| --- | --- | --- | --- | --- |
| Anther | CellStitch* | **0.66 ± 0.10** | **0.53 ± 0.11** | **0.42 ± 0.10** |
|  | Cellpose2D | 0.48 ± 0.11 | **0.53 ± 0.10** | 0.33 ± 0.08 |
|  | Cellpose3D | 0.41 ± 0.17 | 0.38 ± 0.13 | 0.24 ± 0.10 |
| Filament | CellStitch* | **0.74 ± 0.14** | 0.53 ± 0.12 | **0.46 ± 0.13** |
|  | Cellpose2D | 0.51 ± 0.15 | **0.57 ± 0.13** | 0.38 ± 0.13 |
|  | Cellpose3D | 0.03 ± 0.02 | 0.22 ± 0.11 | 0.03 ± 0.02 |
| Leaf | CellStitch* | **0.83 ± 0.17** | **0.75 ± 0.19** | **0.68 ± 0.21** |
|  | Cellpose2D | 0.65 ± 0.19 | **0.77 ± 0.19** | 0.56 ± 0.20 |
|  | Cellpose3D | 0.07 ± 0.05 | 0.53 ± 0.20 | 0.06 ± 0.04 |
| Pedicel | CellStitch* | 0.57 ± 0.23 | **0.44 ± 0.16** | **0.35 ± 0.16** |
|  | Cellpose2D | 0.36 ± 0.19 | **0.44 ± 0.15** | 0.25 ± 0.13 |
|  | Cellpose3D | **0.61 ± 0.25** | 0.30 ± 0.13 | 0.27 ± 0.13 |
| Sepal | CellStitch* | **0.51 ± 0.12** | **0.43 ± 0.16** | **0.31 ± 0.12** |
|  | Cellpose2D | 0.33 ± 0.11 | **0.44 ± 0.17** | 0.23 ± 0.09 |
|  | Cellpose3D | 0.41 ± 0.23 | 0.34 ± 0.12 | 0.20 ± 0.10 |
| Valve | CellStitch* | **0.71 ± 0.07** | 0.47 ± 0.04 | **0.40 ± 0.05** |
|  | Cellpose2D | 0.58 ± 0.09 | 0.47 ± 0.04 | 0.35 ± 0.05 |
|  | Cellpose3D* | 0.67 ± 0.10 | **0.51 ± 0.05** | **0.41 ± 0.06** |

**Figure 6:**
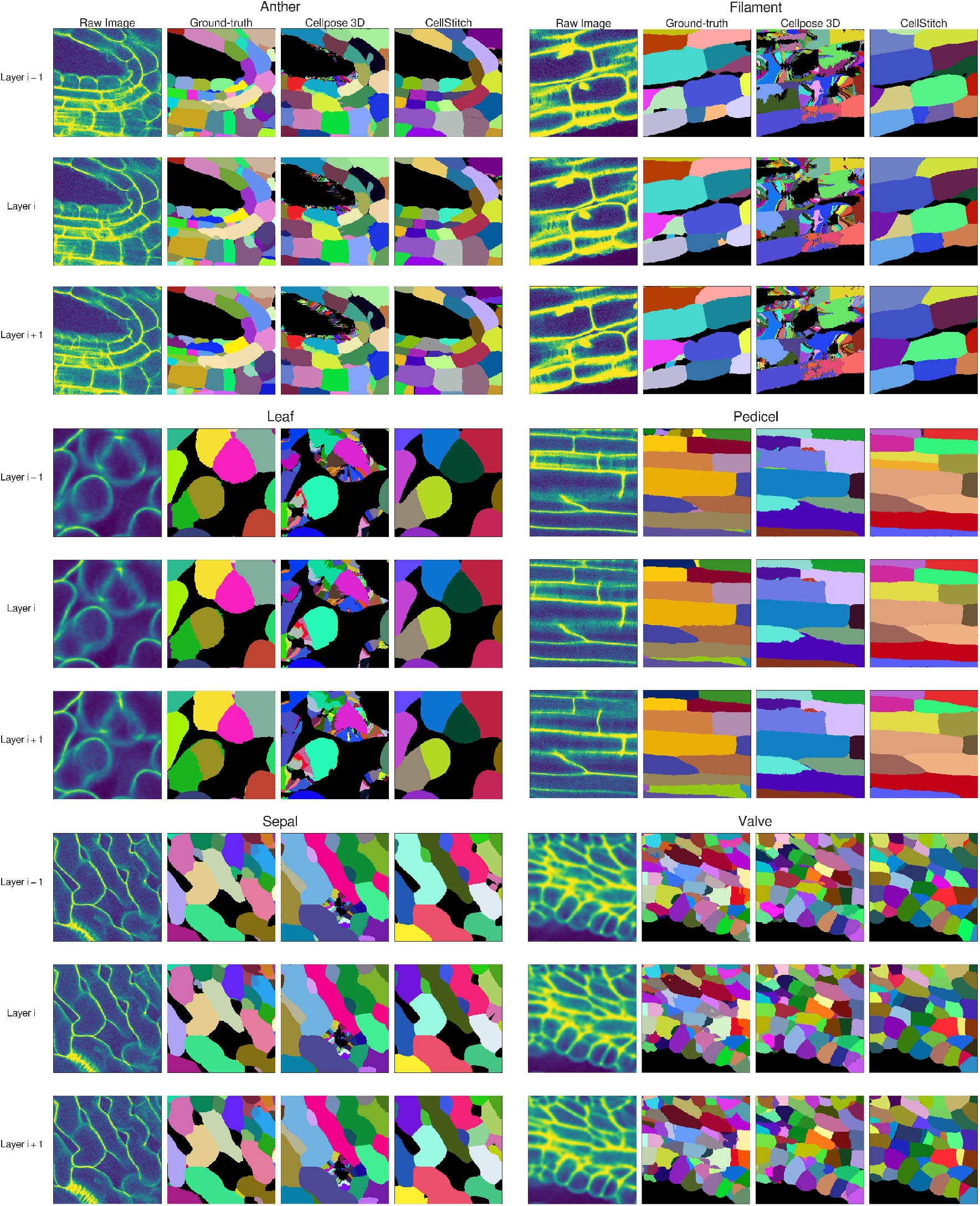
Pipeline results on Arabidopsis 3D Digital Tissue Atlas. Visualization of raw image, ground-truth annotations, Cellpose3D & CellStitch (left to right) on various tissue regions of Arabidopsis.

## 5 Discussion

We developed CellStitch in order to address the challenge of 3D segmentation of anisotropy images. Existing methods fail to incorporate information from all dimensions (in 2D-based methods), and are not capable of generalizing isotropic frameworks to the anisotropic setting (2.5D-based methods) with the lack of training data (3D-based). Our framework relies on optimal transport to align cell slices from different layers in order to reconstruct 3D segmentation from 2D layers. A limitation of CellStitch and the 2D-based approaches in general is the dependence on the chosen 2D segmentation framework; if the 2D segmentations are unreliable, it is unlikely that CellStitch will produce satisfying results. Due to the rapid development of the deep learning-based 2D segmentation pipeline along with increasing 2D training data, we observe that the 2D segmentation by Cellpose [8] achieves desirable 2D masks on the datasets used in this study — nonetheless, future work will aim to extend CellStitch to further improve existing 2D segmentation methods in order to automate 3D segmentation on more complex tissues.

## 6 Conclusions

Recent advances in imaging and sequencing technologies enable *in situ* profiling of cell morphology and gene expression in 3D. Deep learning-based 3D segmentation has achieved great success in biomedical imaging [32, 33]. However, various limitations have hindered the wider applications of 3D segmentation to molecular level data, including fluorescence microscopy imaging.

In this paper, we introduce CellStitch, an efficient algorithm to reconstruct 3D cell instances via layer-wise alignment of 2D segmentation results, optionally followed by an instance-wise interpolation modality for users to recover isotropic cell morphologies from highly anisotropic images. CellStitch bridges the gap between the well-studied 2D segmentation problem and the increasing demand for 3D cell segmentation and extends a flexible pipeline to leverage full 3D instance segmentation. Through previous stitching and pipeline benchmarking, CellStitch demonstrated robustness over other 2D, 2.5D, and 3D methods under various anisotropy levels, which is common among shallow-depth 3D microscopic tissue images. Our interpolation results also provide potential insights for *in situ* experimental designs, where nuclei or cytoplasm signals could be stained in less dense layers than spatial RNA maps, given the cost and technical difficulties to perform multiplexed mRNA, DAPI, and cytoplasm marker imaging.

## Acknowledgments

We would like to thank Itsik Pe’er for many helpful comments based on a careful reading of a previous draft. We further thank Steven Yin, Alex Toberoff, James Wang, members of the Pe’er lab and the Irving Institute for Cancer Dynamics for valuable discussions. Y.J. acknowledges support from the Columbia University Presidential Fellowship. This project has been made possible in part by grant number 2022-253560 from the Chan Zuckerberg Initiative DAF, an advised fund of Silicon Valley Community Foundation, a Columbia University HICCC-SEAS seed grant, DARPA grant HR001119S0076, and ONR grant N00014-22-1-2679.

## Author Contributions

YL: Conceptualization, Methodology, Software, Writing-Original draft preparation. YJ: Conceptualization, Methodology, Software Validation, Visualization, Writing-Original draft preparation. EA: Conceptualization, Writing-Reviewing and Editing, Supervision. AJB: Conceptualization, Writing-Reviewing and Editing, Supervision.

## Availability of data and materials

A python implementation of this method, CellStitch, is available at https://github.com/imyiningliu/cellstitch. The reproduction of all experiments presented herein can be accessed via https://github.com/imyiningliu/cellstitch/tree/main/notebooks.

## Declarations

### Ethics approval and consent to participate

Not applicable.

### Consent for publication

Not applicable.

